# RNA-dependent intergenerational inheritance of enhanced synaptic plasticity after environmental enrichment

**DOI:** 10.1101/178814

**Authors:** Eva Benito, Cemil Kerimoglu, Binu Ramachandran, Qihui Zhou, Tonatiuh Pena, Vincenzo Capece, Gaurav Jain, Susanne Burkhardt, Roman Stilling, Dieter Edbauer, Camin Dean, André Fischer

## Abstract

Physical exercise in combination with cognitive training is known to enhance synaptic plasticity, learning & memory and lower the risk for various complex diseases including Alzheimer’s disease. Here we show that exposure of adult male mice to an environmental enrichment paradigm leads to enhancement of synaptic plasticity and cognition also in the next generation. We show that the effect is mediated through sperm RNA and is explained by microRNAs 212/132. In conclusion, our study reports intergenerational inheritance of an acquired cognitive benefit and points to specific microRNAs as candidates mechanistically involved in this type of transmission.

## INTRODUCTION

There is emerging evidence that exposure to environmental stimuli can initiate processes that transmit information to the next generation via non-genetic mechanisms (1-3). Such forms of inter- or transgenerational inheritance have been described for aversive stimuli, such as chronic or early life stress that lead to altered response of the HPA axis, increased anxiety and depressive-like behavior in the following generations (4) (5). There is also evidence that exposure of individuals to detrimental environmental stimuli can lead to cellular adaptations that protect the offspring when they are exposed to the same environmental insult (6). The idea that environmental factors can affect germ cells and thereby alter biological processes in the offspring is fascinating and may play an important role in the pathogenesis of complex diseases, especially in neuropsychiatric disorders (7) (8).

An environmental factor that was shown to lower the risk for various complex diseases, including those affecting the brain, is the combination of physical exercise and cognitive training, also called environmental enrichment (EE). EE is known to enhance synaptic plasticity in rodents and humans and is thus considered a suitable strategy to reduce the risk for dementia and other cognitive diseases (9) (10) (11) (12). Importantly, there is evidence that exposure of juvenile mice to EE can enhance hippocampal synaptic plasticity in their offspring (13). Whether EE training in adulthood might also affect synaptic function of the next generation has not been tested so far and the underlying mechanisms of transgenerational transmission are still poorly understood. There is however evidence that RNA in gametes could play a role (2, 3).

In this study we demonstrate that exposure of adult mice to EE significantly enhances hippocampal LTP in offspring that did not undergo EE exposure. We show that this effect is mechanistically linked to RNA in sperm and identify microRNAs 212/132 as one of the key factors. In fact, blocking the action of microRNAs 212/132 in oocytes injected with RNA from EE fathers reverses intergenerational LTP enhancement. Finally we show that in addition to LTP, offspring born to EE fathers or those born to fertilized oocytes injected with sperm RNA of EE mice show a cognitive benefit that is however independent of sperm microRNA 212/132.

## RESULTS

### EE training increases hippocampal synaptic plasticity in an intergenerational manner

First, we wanted to confirm that our EE protocol enhances hippocampal LTP in adult mice. To this end we subjected mice to 10 weeks of EE training before measuring hippocampal LTP at the Schaffer Collateral CA1 synapse. We observed a highly significant increase in LTP (Figure 1A). Next, we tested whether EE training in adult male mice would affect synaptic plasticity in their offspring. To this end adult male mice were subjected to 10 weeks of EE training and then mated to home-caged females (Fig 1B). We then measured hippocampal LTP when the offspring were adult (3 months of age). Notably, offspring of EE fathers had increased LTP compared to those born to fathers that were housed in home cages (HC controls; Fig 1C). The effect was similar in both male and female offspring (Fig S1). We then tested whether this phenotype is passed on to the grandoffspring of the original EE animals, representing the F2 generation. We did not observe any difference in the LTP levels in F2 mice compared to controls (Fig 1D). These data indicate that EE in male mice leads to enhanced hippocampal synaptic plasticity in offspring, but that this effect represents inter- (and not trans-) generational inheritance.

**Figure 1.**
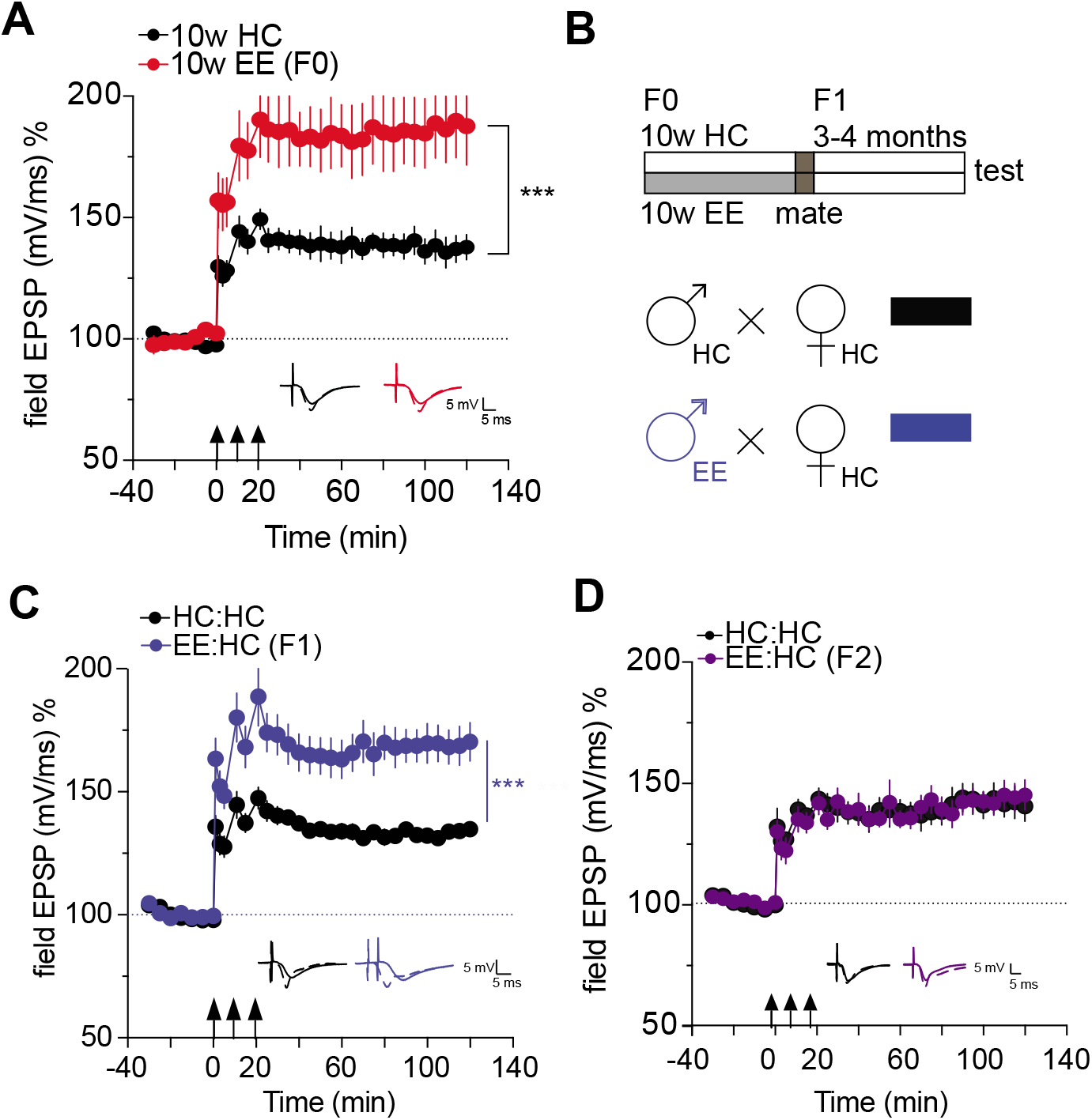
Intergenerational inheritance of enhanced LTP through environmental enrichment. **A.** LTP is enhanced in 10w EE compared to HC controls. *** p < 0.001 for main effect treatment repeated measures ANOVA (F (1, 12) = 11.86); n = 8 (10w EE); 6 (10w HC). **B.** Mating scheme: mice are subjected to EE for 10 weeks, mated and the offspring are tested 3-4 months after birth. Controls spend the same amount of time in the HC. **C.** LTP is enhanced in mice born to EE fathers EE:HC (n = 10) compared to HC:HC (n = 10) controls. p < 0.001 for main effect treatment HC:HC vs. EE:HC, repeated measures ANOVA (F (1, 18) = 18.74). **D.** LTP is not enhanced in the F2 generation. *** pval < 0.001, * pval < 0.05, error bars indicate SEM.

### Intergenerational enhancement of LTP is mediated via sperm RNA

Since the mothers were never exposed to EE, which might have affected maternal care, the enhanced synaptic plasticity observed in offspring of fathers that underwent EE training must be associated with changes in the father’s gametes. Previous reports linked sperm RNA to transgenerational inheritance and observed for example that individual microRNAs were altered in sperm of mice that passed an acquired anxiety phenotype to the next generation (14). In addition, there is evidence that manipulating microRNA levels in gametes can alter the offspring’s phenotype (15) (16).At the same time, microRNAs are known to play key roles in promoting synaptic plasticity (17) (1). We therefore hypothesized that microRNAs might play a role in the intergenerational transmission of EE-induced LTP enhancement. Therefore we measured the sperm microRNAome detectable in mice used in our experimental system and specifically searched the corresponding data for microRNAs that were 1) expressed in sperm, 2) have been linked to brain plasticity and memory function and 3) have a documented role in brain development since this might be a possible route of action by which microRNAs present in gametes could affect brain function (Fig S2).

By this we identified miR-212 and miR-132, which are co-expressed and have been shown to affect synaptic function and learning behavior in mice and play a role in brain development (18) (19) (20-22). Therefore we assayed the expression of the miR212/132 cluster in mice upon EE training. We found that miR132 and miR212 were up-regulated in the hippocampus and in sperm of mice that were exposed to EE for 10 weeks (Fig 2A-B). These findings led us to hypothesize that sperm RNA, and in particular miR212/132 might play a role in the intergenerational inheritance of the EE phenotype. To test this possibility we first injected RNA from sperm of HC or EE fathers into fertilized oocytes and examined LTP in the offspring once they were adult. RNA in both groups was co-injected with scrambled RNA allowing us to include a third group in which we injected into oocytes sperm RNA from EE fathers along with miR-212/132 inhibitor, which we had previously validated for their inhibitory action (Fig S3).

**Figure 2.**
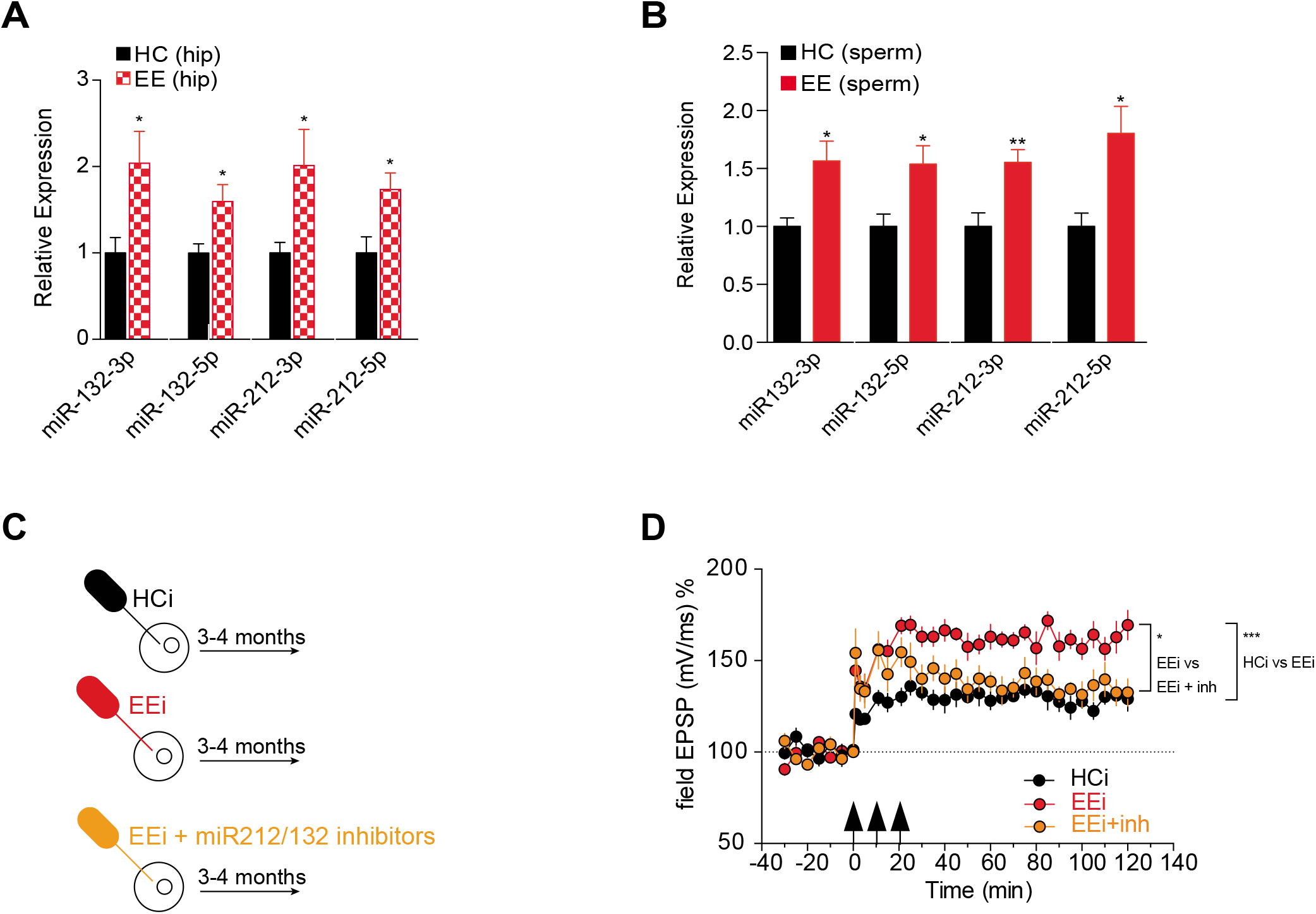
microRNAs 212/132 are increased in brain and sperm of EE males and they are involved in intergenerational inheritance of the enhanced LTP phenotype. **A**. qRT-PCR from hippocampus of 10w EE males demonstrates increased expression of miR212/132 (** *p* < 0.01, * *p* < 0.05). **B.** qRT-PCR from sperm of 10wEE males demonstrates increased expression of miR212/132 (** *p* < 0.01, * *p* < 0.05). **C.** Oocyte injection scheme: controls were injected with sperm RNA from HC fathers + miRNA inhibitor negative control. EEi oocytes were injected with sperm RNA from EE fathers + miRNA inhibitor negative control. EEi + miRNA inhibitors were injected with RNA from EE fathers + miR212/132 inhibitors. **D.** LTP is significantly elevated in EE-sperm-RNA-injected animals compared to HC-sperm-RNA-injected controls, and coinjection with miR-212/132 inhibitors (EEi + inh) reverses the LTP enhancement. *** *p* < 0.001 repeated measures ANOVA, main effect HCi vs. EEi (F (1, 8) = 52.90). * *p* < 0.05 repeated measures ANOVA, main effect EE vs. EE+inh (F (1, 9) = 6.099). n = 5 (HCi); 5 (EEi); 6 (EEi+inhibitors). Error bars indicate SEM.

We observed that the offspring born to oocytes injected with RNA from 10w EE mice exhibited enhanced LTP, which was reversed to control (HC) levels if miR-212/132 inhibitors were co-administered (Fig 2C-D). These data demonstrate that EE in adult males enhances hippocampal synaptic plasticity in offspring and that this effect is mediated through sperm RNA causally involving miR212/132.

Next, we wondered whether this enhancement of LTP was accompanied by an improvement in cognitive performance. To this end, offspring of EE fathers and those of HC controls were subjected to behavior testing (Fig 3A). We observed no difference in explorative behavior and basal anxiety amongst groups (Fig S4). Next mice were subjected to two hippocampus-dependent behavioral tests, namely the contextual pavlovian fear conditioning (FC) and Morris Water Maze (MWM) paradigms. To avoid ceiling effects on learning, mice were trained using rather mild protocols. Thus, the electric footshock applied during fear conditioning was 0.5mA, an intensity that allows the detection of memory improvement. In the MWM, mice were trained for a maximum of 5 days, a protocol that in our hands results in a moderate memory consolidation of the platform location in WT animals (Fig S5), that normally form robust spatial memory only after 8 training days (Fig S5).

**Figure 3.**
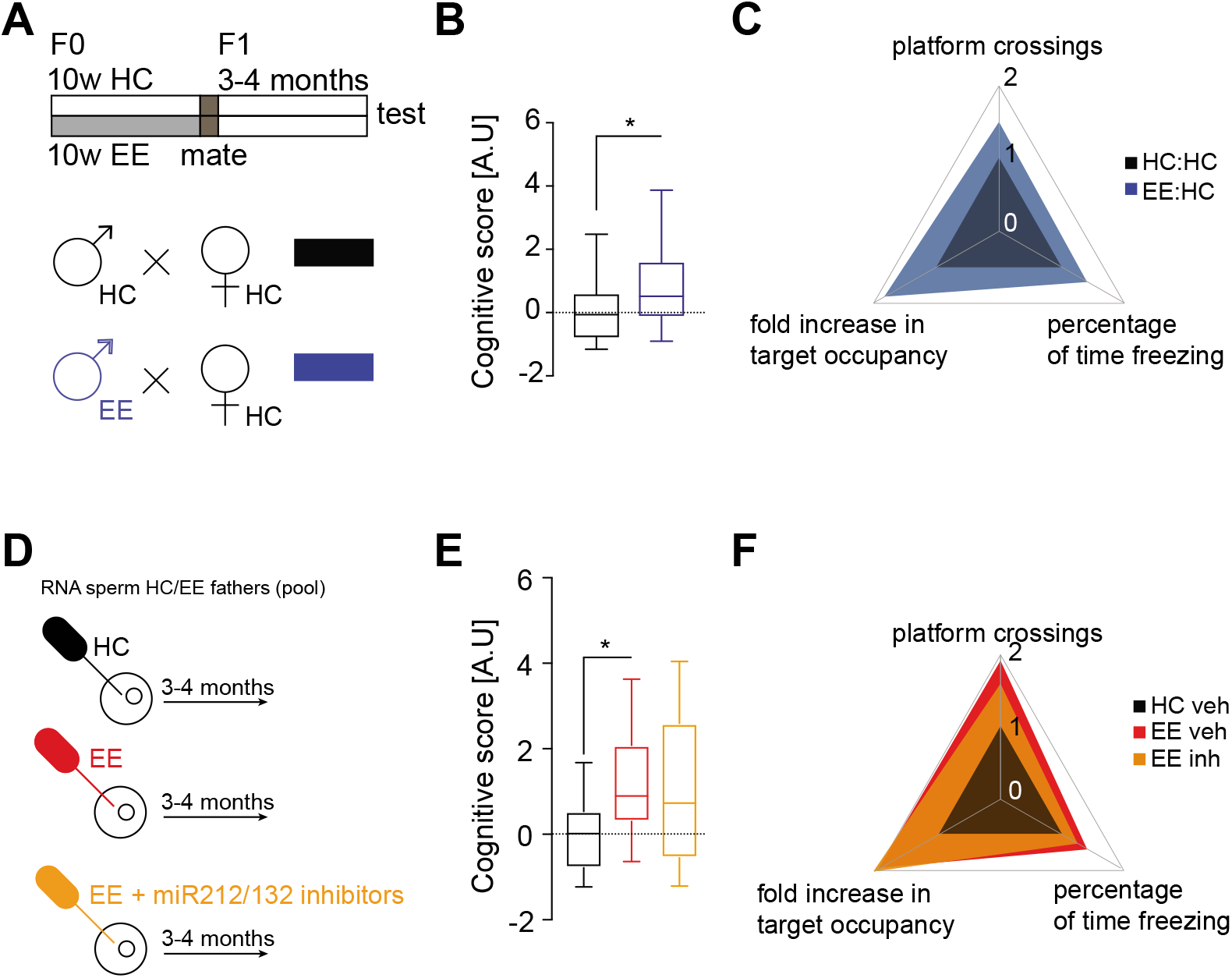
Mice born to EE fathers have a mild but significant cognitive advantage. **A.** Breeding scheme. **B.** Mice born to EE fathers have a significantly bigger cognitive score. **C.** Plot illustrating the magnitude of change of each individual parameter that went into the cognitive score calculation. **D.** Oocyte injection scheme. Sperm RNA from a pool of HC or EE fathers was isolated and mixed with scrambled negative control or miR212/132 inhibitors and injected into the cytoplasm of fertilized oocytes. The offspring from these injections were then tested in several behavioral tasks at the age of 3-4 months. **E.** Injection of EE sperm RNA into fertilized oocytes provides a cognitive advantage to the offspring, as reflected in the significant increase in the cognitive score. **F.** Plot illustrating the magnitude of change in the different groups of each individual parameter of the cognitive score. Significance for the F1 generation was calculated using linear mixed models to account for batch and litter effects (see methods). * *p* < 0.05 (t-value = 2.80, Df = 10). Significance for the offspring of oocyte injections was calculated using a two-tailed Student’s *t*-test (see methods). * *p* < 0.05. n = 29, N = 6 (HC:HC), n = 32, N = 7 (EE:HC), n = 9 (HC sperm RNA + negative control oocyte injections), n = 14 (EE sperm RNA + negative control oocyte injections), n = 18 (EE sperm RNA + miR-212/132 inhibitor oocyte injections).

First we tested mice in the contextual fear conditioning. We observed that mice born to EE fathers showed elevated freezing behavior when compared to mice born to HC fathers (Fig S6A, *t*-test, P< 0.05 for HC:HC vs. EE:HC group). Similar results were observed in the water maze paradigm (Fig S6C-D, *t*-test P< 0.05, HC:HC vs EE:HC group). We also employed a more stringent approach using a linear-mixed model for statistical analysis taking into account the effect of the different litters, since for the behavioral experiments we employed more than one mouse/litter for the analysis. In this analysis we observed a non-significant trend for memory enhancement in offspring from EE fathers in the fear conditioning and water maze paradigm (percentage of time freezing: t = 1.45, Df = 10, pval = 0.17; platform crossings: t = 1.79, Df = 10, pval = 0.1; relative time of platform occupancy: t = 1.99, Df = 10, pval = 0.07). In sum these data suggest that the intergenerational effect of EE on memory function is more subtle compared to the LTP phenotype. This may in part be due to the fact that the analysis of LTP in hippocampal slice preparation allows for a well-defined and specific readout while the analysis of an animal’s complex behavior as an estimate of memory function, on the other hand, offers a rather limited dynamic range. Rather than relying on the classical readout of the fear conditioning and water maze paradigms we therefore decided to also calculate an integrated cognitive score based on a principal component analysis (PCA). The first component of the PCA analysis captures the most variance and hence we took the scores from the PC1 (principal component 1) as a “cognitive score” that would reflect the overall change in cognitive function. This score revealed a significant difference between animals born to a HC or an EE father (Fig 3B-C linear mixed model t-value = 2.80, Df = 10, pval = 0. 018), confirming that there is indeed intergenerational inheritance of a mild, but significant cognitive advantage after EE exposure in the fathers.

Next we decided to address the question of whether sperm RNA and in particular microRNA212/132 would pay a role in the memory enhancement seen in offspring born to EE fathers. As described for the LTP experiments (See Fig 2) we isolated RNA from sperm of EE fathers and injected this RNA into fertilized oocytes with scrambled RNA. Oocytes injected with RNA from sperm of HC fathers together with scrambled RNA were used as control. Also here, we included a group where EE sperm RNA was co-injected with miR212/132 inhibitors.

The resulting mice were subjected to behavior testing when they were adult (Fig 3D). Mice that originated from oocytes injected with sperm RNA from EE fathers showed again improved memory function in the fear conditioning (Fig S6F, pval = 0.04, *t*-test) and water maze paradigms (Fig S5G-H, platform crossings: pval =0.01, Mann-Whitney test; relative time of platform occupancy: pval =, 0.04, *t*-test.) when compared to mice that originated from oocytes injected with sperm RNA from non-EE fathers (HC), confirming our previous results on LTP enhancement. This observation was further supported when we analyzed the cognitive score as described above (Fig 3E-F), suggesting that similar to the intergenerational inheritance of cognitive enhancement mediated by EE, injection of EE sperm RNA into fertilized oocytes also results in a cognitive benefit. In contrast to the effect on LTP the miR212/132 cluster appeared to have no effect on the behavioral readout. Mice that were born from oocytes injected with sperm RNA from EE fathers and co-injected with miR212/132 inhibitors exhibited a similar, albeit insignificant, trend for memory enhancement (Fig 3E and Fig S6F-H; percentage time freezing: pval = 0.28, Mann-Whitney test; platform crossings: pval = 0.19, Mann-Whitney test; fold increase in target occupancy: pval = 0.08, *t*-test). These findings suggest that the intergenerational effect of EE on LTP and memory enhancement critically depends on sperm RNA and that the LTP effect is mediated via altered levels of miR212/132. In contrast, miR212/132 levels cannot explain the enhanced memory function indicating that additional mechanisms contribute to the intergenerational enhancement of learning behavior in response to EE.

## DISCUSSION

Our data demonstrate that EE during adulthood mediates the enhancement of hippocampal LTP in the adult offspring. This is not only the first confirmation but also an important extension to the original observation by Arai et al. (13) who showed that EE in juvenile mice (2 weeks of age) enhances LTP in offspring. Our observation that this phenomenon occurs even when EE is initiated at a time point at which brain development is complete has important implications, suggesting that exercise before conception could provide a brain plasticity benefit to the offspring. Moreover, we provide evidence that in addition to enhanced LTP offspring of EE fathers exhibit a mild but significant cognitive advantage when compared to offspring of HC fathers. When compared to the LTP effect, the memory enhancement was however moderate. One explanation could be that the analysis of Schaffer collateral CA1 LTP in a hippocampal slice preparation is much more sensitive and thus better suited to detect increased or decreased plasticity when compared to behavioral assays that offer a limited dynamic range for the detection of changes. In line with this, calculation of a combined cognitive score on the basis of two hippocampal memory tests using principal component analysis showed that offspring born to EE fathers or from oocytes treated with RNA isolated from sperm of EE mice exhibit a significant cognitive benefit. It also has to be considered that in our study we analyzed memory function in healthy wild type mice. It will thus be interesting to see if EE training in fathers would provide a benefit in synaptic plasticity and memory function to offspring in a situation when brain plasticity is challenged as it is for example the case during aging or in neurodegenerative conditions. In support of this view recent data shows that EE partially ameliorates the detrimental transgenerational effects observed in offspring born to parents that were exposed to stressful experiences (23). Similarly, it will be interesting to see if increased brain plasticity is observed in offspring when the mother undergoes EE training. In this study we did not address this issue and rather focused on EE fathers since the analysis of the corresponding gametes for subsequent analysis is more suited for an initial study. Another advantage is that EE fathers did not contribute to parental care and since the mothers did not undergo EE, we can exclude that parental care has any influence on the phenotypes reported in our study. It will be interesting to test if EE training in adult female mice will also transmit a synaptic plasticity and cognitive benefit to offspring and to elucidate the underlying mechanisms. Such studies however need to involve cross-fostering experiments to dissociate the effect of EE on maternal care form gamete-specific changes, and thus – at least in our hands – appear more complicated since the cross-fostering procedure itself seems to affect brain plasticity making the interpretation of the data more difficult. Future experiments will address these issues.

Our data show that the intergenerational inheritance of the EE-phenotype involves changes in sperm RNA. This finding is in line with previous studies that investigated transgenerational effects in mutant mice (15), in mice exposed to specific stressors leading to altered glucocorticoid signaling (5, 16), anxiety behavior (14) or diet-induced obesity (24). An important difference to most of these studies is that we observe and inter-but not a transgenerational effect. Thus, an EE-induced brain plasticity benefit was detectable in the F1 but not in the F2 generation. From an evolutionary point of view our finding could make sense, since nature offers to the organism a physiological system that allows for non-genetic inheritance of a cognitive benefit in situations of demand but makes sure that these phenotypes do not persist when the environmental settings change. Taking into account that “too much” plasticity and the resulting aberrant neuronal activity have been linked to neurodegenerative diseases (25, 26) (27) it appears logical that EE mediated LTP enhancement is restricted only to the next generation. Moreover, the EE-mediated intergenerational changes of brain function appear to be regulated at multiple levels. Our data clearly show that one of these mechanisms is related to sperm RNA. In this context it is interesting to note that the EE-mediated inter-generational inheritance of LTP enhancement was linked to the action of miR212/132. To the best of our knowledge this is the first report that inhibition of a specific microRNA cluster in gametes can affect inter-generational phenotypes. These data are also in line with an earlier study that demonstrated that oocyte injection of a mixture of eight microRNAs could recapitulate the transgenerational effects observed in response to chronic paternal stress (16). However, inhibition of miR212/132 in fertilized oocytes injected with sperm RNA from EE fathers did not occlude the inheritance of improved memory function assayed in the behavior paradigms. These data clearly indicate that there must be additional RNA-dependent mechanisms of similar importance. Moreover, we cannot exclude that non-RNA mediated processes also contribute to this phenotype. In fact, altered DNA methylation (28, 29), and histone modifications or the replacement of canonical histones with histone variants (30, 31) have been implicated in intergenerational inheritance. These findings suggest that the intergenerational effects of EE-mediated LTP improvement can be dissociated from the improvement of learning behavior at the molecular level and are in line with the view that although LTP has been considered a molecular correlate of memory function, it cannot fully explain all aspects of memory consolidation (32).

The precise mechanisms by which sperm RNA transmit EE-induced intergenerational enhancement of brain function to adult offspring remain to be identified. One possibility is that sperm microRNAs alter gene expression during embryonic development thereby causing subtle changes in brain plasticity. This hypothesis appears worthwhile to be addressed in future studies, since even small changes in brain development can affect synaptic function in adult organisms, and are believed to contribute – for example – to neuropsychiatric diseases (33, 34). Of note, miR-212/132 have been linked to developmental brain diseases such as schizophrenia and play a role in the developing, as well as in the adult brain (35, 36). These data may also help to understand why EE-induced transmission of LTP enhancement is only observed in the F1 but not in the F2 generation. We speculate that the mechanisms underlying enhanced brain plasticity in the parents and the offspring are different, a hypothesis to be tested in future studies. In support of this view, we observed that hippocampal levels of microRNA132/212 increase in fathers exposed to EE (F0 generation; see Fig 2), a finding that is in line with previous data linking increased microRNA132/212 levels to memory enhancement (37) (21, 22), but not in their adult offspring ( F1 generation; Fig S7).

In conclusion, the idea that EE training in adulthood elicits a cognitive benefit not only for the individual undergoing this procedure but also for its offspring is fascinating. Whether these findings are translatable to humans needs to be determined. Nevertheless, the accumulating evidence that sperm RNA content encodes information about environmentally-induced phenotypic traits is an issue that needs to be considered not only in reproductive medicine, but may also offer the chance to discover new biomarkers for complex diseases.

## MATERIALS AND METHODS

### Animals

All procedures were performed according to protocols approved by the Lower Saxony State Office for Consumer Protection and Food Safety. Wildtype C57Bl/6J animals were ordered from Janvier labs at 9 weeks of age and were allowed to habituate to the animal facility and the group for one week. Afterwards they were housed either in standard home cages (HC) or under environmental enrichment (EE) conditions as described below. Food and water were provided *ad libitum*. Animals were kept on a 12h/12h light/dark cycle. For all experiments adult mice were used which refers to an age of 3-5 month.

### Environmental enrichment

For environmental enrichment, mice were kept in groups of 4-5 in large cages and provided with toys (in the form of tunnels, housing and differently shaped objects, 8 per cage) and running wheels (2 per cage). 2 toys were replaced daily with novel ones and the rest was rearranged inside the cage. Cages were changed weekly altogether. Mice were put in the EE at 10 weeks of age and kept there for 10 weeks. HC animals were also kept in groups of 4-5 in large cages in the absence of objects and running wheels but subjected to the same cleaning schedule.

### Luciferase assay

For luciferase assays, the complementary seed sequence for miR-212/132 was cloned into pmirGLO (Promega) into PmeI/XbaI sites by oligo hybridization (see below). The constructs were cotransfected into HEK293 cells with miR-212/132 mimics (5nM, Qiagen) or miR-212/132 inhibitors (5nM for 3p arms, 50nM for 5p arms, Exiqon) using lipofectamine according to the manufacturer’s instructions. Luciferase measurements were taken 48h in a Tecan plate reader using Promega’s Dual-Luciferase Reporter Assay System following manufacturer’s instructions. The Firefly/Renilla ratio was calculated and then normalized to control conditions.

### Oligonucleotide sequences used for cloning

The following oligonucleotides were used for cloning the miRNA target sequences into pmirGLO:

> miR-132-3p F: AAACTAGCGGCCGCTAGTCGACCATGGCTGTAGACTGTTA
>
> miR-132-3p R: CTAGATAACAGTCTACAGCCATGGTCGACTAGCGGCCGCTAGTTT
>
> miR-132-5p F: AAACTAGCGGCCGCTAGTGTAACAATCGAAAGCCACGGTTT
>
> miR-132-5p R: CTAGAAACCGTGGCTTTCGATTGTTACACTAGCGGCCGCTAGTTT
>
> miR-212-3p F: AAACTAGCGGCCGCTAGTTGGCCGTGACTGGAGACTGTTAT
>
> miR-212-3p R: CTAGATAACAGTCTCCAGTCACGGCCAACTAGCGGCCGCTAGTTT
>
> miR-212-5p F: AAACTAGCGGCCGCTAGTAGTAAGCAGTCTAGAGCCAAGGT

### Sperm collection and RNA isolation

Animals were sacrificed by cervical dislocation. Sperm was isolated by dissecting the epididymis and running a needle through the tube to allow sperm to swim out into 1ml of pre-warmed PBS. Epidydimal tissue was kept at 37^o^C for 20 minutes, the supernatant was then collected and centrifuged at 8000 x g for 5min at 4^o^C. In order to remove contaminating epithelial cells prior to RNA isolation, the pellet was treated with hypotonic buffer (0.1 % SDS, 0.5 % Triton X) for 30 minutes on ice to ensure that endothelial cells from the surrounding tissue were disrupted. The solution was centrifuged once more at 8000 x g for 10 min at 4^o^C to obtain the final sample. RNA was isolated using Tri-Reagent from Sigma-Aldrich according to manufacturer’s instructions. RNA was treated with DNAse I (ThermoFisher) and further purified using phenol:chloroform to remove all DNAse I components. RNA concentration was determined with the Nanodrop and quality with the Bioanalizer.

### Quantitative RT-PCR

cDNA synthesis was done using Qiagen’s Reverse Transcription kit II according to manufacturer’s instructions starting from 250ng of DNAse I-treated RNA. microRNA levels were determined using Qiagen’s miRNA assays. miRNA levels were normalized to RNU6B. 2-5ng of cDNA were used per reaction. Assays were run in duplicate. The ΔΔCt method was used to calculate relative expression.

### RNA injections into fertilized oocytes

Sperm RNA from 4 HC and 4 EE animals was isolated as described above. RNAs were pooled and set at 0.5ng/μl in 250μl 0.1mM EDTA, 5mM pH 7.4 based on a previous publication reporting significant effects at this concentration [Gapp et al. 2014]. 3-4 week old C57Bl/6JRj females were super-ovulated with pregnant male serum gonadotropin (7.5 U, Intervet) and human chorionic gonadotropin (7.5 U, Intervet) and mated with C57Bl/6JRj males. Donor females were sacrificed by cervical dislocation on the day of plug and fertilized eggs were collected. RNA was injected into the cytoplasm of zygotes at the pronuclear stage using an Eppendorf Femtojet and Femtotip II capillaries with constant flow under visual control on an inverted microscope using a 40x air objective and DIC optics. Injected zygotes were transferred into the oviduct of pseudo-pregnant NMRI fosters bilaterally. The offspring generated from these injections was allowed to grow in a standard cage until 3 month of age before behavioral testing.

### Behavioral experiments

Animals that went into behavioral experiments were single-caged 1 week prior to the start of the procedures. For contextual fear conditioning, animals were put in a fear conditioning box (Med Associates) for 3 minutes, after which they received a 2- second 0.5mA footshock. They were left in the box for 10 seconds and then removed and put back to their original cage. 24h later, animals were reintroduced into the conditioning box and freezing behavior was recorded for the 3 minute period. Freezing was considered when animals remained immobile (except for respiratory movements) for at least 2 seconds. Note that this fear conditioning paradigm represents a rather mild training therefore allowing the detection of memory enhancement, which is in line with HC mice showing relatively low freezing levels. For Morris Water Maze, animals were trained in a round pool filled with opaque water where the escape platform was placed approximately 1 cm below water level. They were trained in 4 consecutive 1-minute trials per day with randomized entry points. If the animals did not reach the platform within 1 minute, they were gently guided to it. Animals were allowed to stay on the platform for 15 seconds after each trial. The time needed to reach the platform (escape latency) was automatically recorded via a top-installed camera (TSE) and registered with the TSE software VideoMot. For the probe test, the platform was removed and animals were introduced in the position opposite of the original platform location and left to navigate the pool for 1 minute. The percentage of time spent in the specific region (target time occupancy) where the platform used to be during training was recorded with the TSE software.

### Breeding scheme for intergenerational analysis

10 week-old male mice acquired from Janvier were subjected to 10 weeks of EE according to the protocol described above. HC and EE male mice were bred with females that had remained home-caged throughout this time. Offspring that originated from these pairings were housed in standard home cages and subjected to LTP measurements or behavioral experiments at 3-4 months of age.

### LTP recordings

Acute hippocampal slices were prepared from 3-4 month old mice. Animals were anesthetized with isoflourane and decapitated. Brain was removed from the skull and hippocampus was dissected. Transversal hippocampal slices (400 μm thick) were obtained using a tissue chopper (Stoelting). Slices were collected in ice-cold artificial cerebrospinal fluid (ACSF) (124 mM NaCl, 4.9 mM KCl, 1.2 mM KH2PO4, 2.0 mM MgSO4, 2.0 mM CaCl2, 24.6 mM NaHCO3, 10.0 mM D-glucose; saturated with 95% O2 and 5% CO2; pH 7.4 and 305 mOsm). Slices were incubated in an interface chamber at 32C and high oxygen tension was maintained by bubbling with 95% O2 and 5% CO2 (30 l/h). Slices were allowed to recover for 3 h after preparation. Then monopolar platinum-iridium electrodes (13303, MicroProbes) used for both recording and stimulating were positioned in the CA1 region. The field excitatory postsynaptic potential (fEPSP) slope was recorded with a Model 1700 differential AC amplifier (A-M Systems) and Power 1401 analog-to-digital converter (Cambridge Electronic Design), and monitored on-line with custom-made software, PWIN (IFN Magdeburg). The test stimulation strength was determined for each input as the current needed to elicit a field EPSP of 40% maximal slope. Baseline recording began at least 3.30 h after slice preparation, using test stimuli consisting of four biphasic constant current pulses (0.2 Hz, 0.1 ms/polarity, averaged) per time point, every 5 min for a minimum of 30 min. LTP was induced with a strong tetanization protocol consisting of three stimulus trains (100 biphasic constant-current pulses per train at 100 Hz and 0.2ms/polarity, inter-train interval 10 min). Test stimuli were delivered 1,3,5,11,15,21,25 and 30 min after the first tetanization train and then every 5 min for up to 2 h.

### Statistical analysis

Since several animals from the same litter were analyzed and tests were done at two different time points (i.e. batches) to ensure that the group size was homogeneous and stayed within the same light/dark cycle, we used generalized mixed models to account for such type of random effects. We used the R packages MASS (MASS_7.3-45) and nlme (lme4_1.1-12) to construct and evaluate the generalized linear models. We first constructed a model with treatment and gender as fixed factors and litter nested within batch as random factors. There was no significance for gender main effect or for the interaction between gender and treatment, so we dropped the gender effect for evaluating the main effect of treatment. For the oocyte injection experiments the sperm RNA comes from a pool of 4 HC/EE mice and the manipulation happens at the level of each individual embryo so we considered animals separately. Other analyses were done with GraphPad Prism. All figures represent mean +/-SEM. Exact n (mouse) and N (litter) numbers are specified in figures legends if applicable.

## Acknowledgments

We thank Ursula Fünfschilling from the transgenenic animal facility of the Max Planck Institute for Experimental Medicine, Göttingen, Germany for help with oocyte injections. This work was supported by the following third party funds to AF: DFG project 179/1-1/2013, the DFG priority program 1738, an ERC consolidator grant and funds from the German Center for Neurodegenerative Diseases (DZNE). CD was supported by the European Research Council (FP7/260916), Alexander von Humboldt-Stiftung (Sofja Kovalevskaja), and DFG (DE1951 and SFB889).

## Author contribution

BR, QZ, VC, SB, RS, DE, CD performed experiments and analyzed data. EB, CK and AF designed and performed experiments, analyzed data and wrote the manuscript.

